# Bark traits affect epiphytic bryophyte community assembly in a temperate forest

**DOI:** 10.1101/2023.07.12.548789

**Authors:** Shinichi Tatsumi, Takayuki Ohgue, Wakana A. Azuma, Keita Nishizawa

## Abstract

Bark traits of trees often serve as a key factor determining the community structure of epiphytes. However, the extent to which barks modulate the relative importance of abiotic and biotic assembly processes of epiphytes is poorly understood. Here, using a community phylogenetic approach, we aimed to infer the assembly processes of epiphytic mosses and liverworts on tree species with varying bark traits in a temperate forest of central Japan. We observed a total of 56 moss and 35 liverwort species on 150 trees. Moss communities showed decreasing species richness and a tendency toward phylogenetic overdispersion, that is, higher phylogenetic diversity than expected by chance, in relation to increasing bark roughness, acidity, and wetness. Along the same bark gradients, liverwort communities became phylogenetically clustered. Species richness of both mosses and liverworts increased with the nitrogen content of barks. The results indicate non-random assembly processes such as interspecific competition on resource-rich barks and abiotic filtering associated with environmental harshness and microhabitat variety determined by barks. Our findings imply that bark traits modulate community assembly processes through which epiphyte diversity is maintained.

## Introduction

Epiphytes are essential components of forest biodiversity (Burns and Zotz 2010; Mendieta-Leiva and Zotz 2015; Tatsumi et al. 2017). Understanding the processes through which epiphyte species assemble on host trees provides a crucial step toward developing effective conservation strategies and preserving the functional roles epiphytes play in forest ecosystems (Ellis 2012). Notably, the characteristics of barks have been recognized as a key determinant of epiphyte community structure (Callaway et al. 2002; Wyse and Burns 2011; Mendieta-Leiva and Zotz 2015). However, despite extensive research describing the composition and distribution patterns of epiphytes on various barks, comparatively little is known about how bark traits modulate the relative importance of assembly processes (e.g., abiotic filtering or biotic interactions) driving such patterns (Spicer and Woods 2022).

Phylogenetic diversity has been widely employed to account for evolutionary and ecological relatedness among species within a community. In particular, the sign and magnitude of phylogenetic diversity deviating from null expectations have commonly served as proxies representing the relative strengths of different assembly processes (Webb et al. 2002; Cavender-Bares et al. 2004; Gerhold et al. 2015). Under evolutionary niche conservatism, phylogenetic diversity lower or higher than expected by chance, referred to as phylogenetic clustering and overdispersion, respectively, has been interpreted as indicative of abiotic and biotic assembly (Webb 2000; Webb et al. 2002). In combination with demonstrable environmental gradients, phylogenetic diversity can provide insights into ecological processes through which species assemble into communities (Cadotte and Tucker 2017; Cadotte et al. 2019; Tatsumi et al. 2019).

Here, we explore community assembly of epiphytic bryophytes on barks. Specifically, we analyze phylogenetic diversity of mosses and liverworts, which constitute two major clades of bryophytes, on multiple tree species that represent gradients of bark traits in a temperate forest. Using null models, we test whether communities show tendency toward phylogenetic clustering or overdispersion along the gradients. Based on the phylogenetic community structure observed, we infer underlying assembly processes and their links to bark traits.

## Methods

### Study site and tree species

This study was conducted in the Ashiu Forest Research Station of Kyoto University, western Japan (4186 ha; 35.3° N, 135.8° E; 355 to 959 m elevation) (Fig. S1). The study site is covered by pristine forests and is designated as a National Bryophyte Heritage Site of Japan for its rich bryophyte flora. The mean monthly temperature ranges from −0.4°C in January to 24.0°C in August. The mean annual precipitation is 2568 mm.

We selected 10 tree species for our study: *Acer pictum* subsp. *mono, Acer sieboldianum, Aesculus turbinata, Betula grossa, Castanea crenata, Clethra barbinervis, Cryptomeria japonica, Fagus crenata, Quercus crispula*, and *Quercus serrata*. These species were selected to cover a wide range of bark traits as possible. For each tree species, we surveyed bryophyte communities on 15 trees, totalling 150 trees, in six plots distributed across the study area (Fig. S1). The surveyed were selected in such a way that all tree species had similar levels of variation in tree sizes (Fig. S2) and among-individual geographical distances (Fig. S1).

### Bryophyte survey and diversity

In October 2016, we surveyed epiphytic bryophytes in four 10-cm wide, 200-cm high quadrats positioned at the cardinal directions of each tree, totalling 8000 cm^2^ per tree. We recorded the presence or absence of bryophyte species on each tree. Species were identified in the field or in the laboratory under a microscope. To prevent epigeic species from being included, the quadrats were placed approximately 5–30 cm above the ground surface, depending on the inclination of stems and slopes. We used quadrats with a fixed size so that bryophyte diversity would be comparable among trees of different sizes, without being affected by variation in the survey area *per se*. All trees were surveyed at their cardinal directions to keep the possible influences of aspect consistent.

A bryophyte phylogeny was reconstructed based on three chloroplast genes (*rbc*L, *rps*4, and *trn*L-F), which are commonly used in bryophyte phylogenetics (Stech and Quandt 2010). See Supplementary text 1 for details on phylogeny reconstruction.

We quantified phylogenetic diversity of bryophyte communities using mean pairwise distance (MPD) (Webb 2000). We calculated the standardized effect size of MPD, referred to as net relatedness index (NRI), based on null modelling (Webb et al. 2002). The NRI was defined as (x-μ_null_)/σ_null_, where x is the observed MPD, μ_null_ is the mean MPD of a null distribution, and σ_null_ is the standard deviation of a null distribution (Webb et al. 2002). The null distributions were generated based on 999 iterations of presence-absence randomizations across 150 communities using the independent swap algorithm (Gotelli 2000). Randomizations were conducted separately for mosses and liverworts.

### Bark traits

For each of the 10 tree species, we measured bark roughness, water content, pH, and inorganic nitrogen content. These traits were selected based on previous findings on their associations with epiphyte community structure (Gustafsson and Eriksson, 1995; reviewed by Ellis, 2012). See Supplementary text 2 for details of the measurement methods and Table S1 for the observed bark trait values. We conducted principal component analysis to obtain composite measures of bark traits.

### Regression analyses

We tested the changes in bryophyte species richness along bark trait gradients using generalized linear mixed models with a Poisson error distribution and a log-link function. Changes in MPD were tested using log-normal linear mixed models. Changes in NRI were tested using linear mixed models. We included ‘plots’ as a random variable in all models. We used R 4.3.0 (R Core Team 2023) for all statistical analyses.

## Results and Discussion

We observed a total of 56 moss and 35 liverwort species on 150 trees, with 1016 occurrences of mosses and 515 occurrences of liverworts. Regarding bark traits, more than half of the variation was captured by the first principal component (PC 1) (Fig. 1). The PC 1 represented a composite gradient of bark roughness, pH, and water content, along which we found significant changes in species richness of mosses (Fig. 2a). This result may reflect the impact of bark acidity (ranging from pH 4.16 to 6.18; Table S1), which often reduce germination and growth rates of mosses (Löbel and Rydin 2010), thereby leading to a decrease in species richness (Kaufmann et al. 2019; Mitchell et al. 2021).

**Figure 1.**
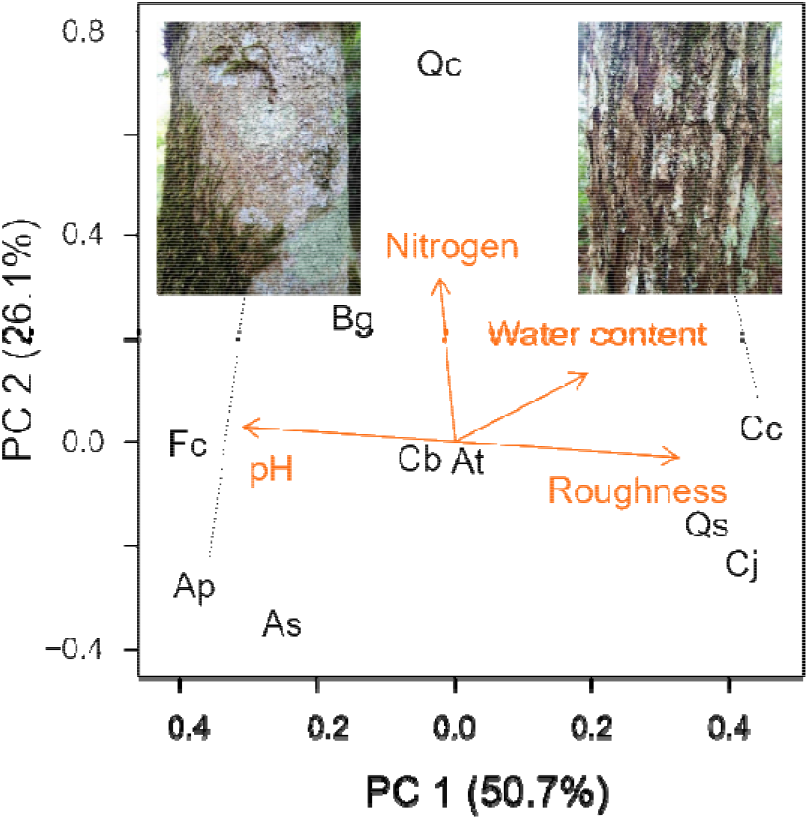
Ordination plot of tree bark traits. Values in parentheses indicate the proportion of variation explained by the first and second principal components (PC). Ap = *Acer pictum* subsp. *mono*, As = *Acer sieboldianum*, At = *Aesculus turbinata*, Bg = *Betula grossa*, Cc = *Castanea crenata*, Cb = *Clethra barbinervis*, Cj = *Cryptomeria japonica*, Fc = *Fagus crenata*, Qc = *Quercus crispula*, and Qs = *Quercus serrata*. The pictures show barks of Ap and Cc as examples with contrasting barks.

**Figure 2.**
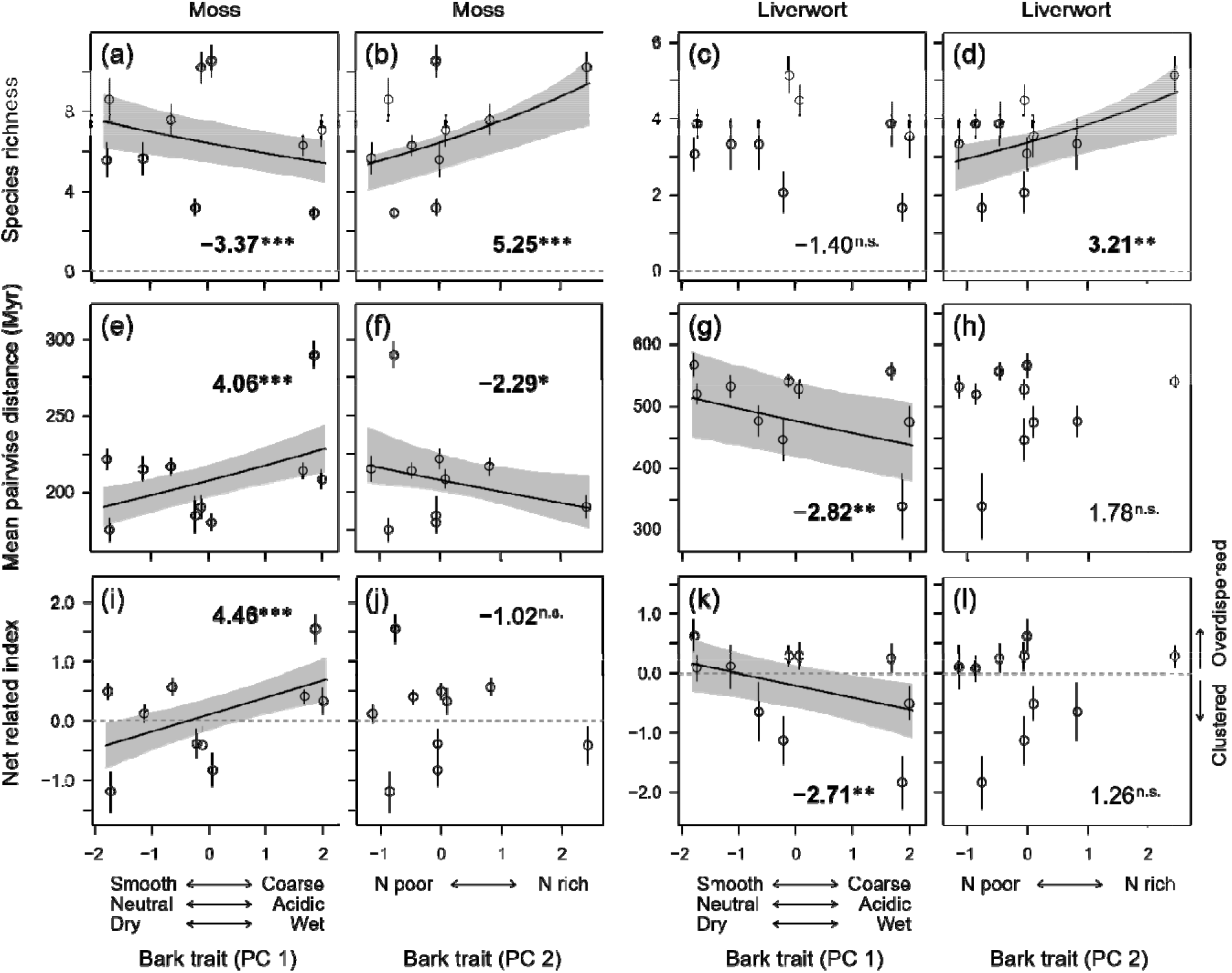
Changes in bryophyte diversity along bark trait gradients. **(a, b, c, d)** Species richness, **(e, f, g, h)** mean phylogenetic diversity, and **(i, j, k, l)** net relatedness indices of mosses and liverworts along the first and second principal components (PC 1 and PC 2) of bark traits. Circles and vertical bars represent the mean and standard errors for each tree species (*n* = 15 trees surveyed for each of ten tree species, totalling *n* = 150). Lines show fitted models with significant slopes (*P* < 0.05). Grey areas represent 95% confidence intervals of the fitted models. The values in each panel indicate the z-statistics of the slope of the fitted model. Significance: *, *P* < 0.05; **, *P* < 0.01; ***, *P* < 0.001; not significant (n.s.), *P* ≥ 0.05.

The MPD and NRI of mosses increased along the PC 1 axis (Fig. 2e, 2i), suggesting changes in assembly processes. Specifically, moss communities became phylogenetically overdispersed (NRI > 0) on rough, acid, and wet barks (Fig. 2i); that is, communities became composed of species belonging to a larger variety of lineages than would be expected by chance. A possible reason for this pattern is that rough barks, which often have a greater heterogeneity of microhabitats than smooth barks (Wyse and Burns 2011; Lamit et al. 2015), allowed moss species from different lineages favouring different microhabitats to coexist. Alternatively, the observed pattern of overdispersion may reflect independent adaptations among moss lineages to harsh environments.

Convergent evolution of plants to harsh environments is a commonly observed phenomenon, including adaptations of alpine plants to high elevations (Bryant et al. 2008) and mangrove trees to salinity (Shi et al. 2005). In our study, acidity may have acted as an environmental filter representing harshness, given the fact that many extant moss species favour neutral pH (Robinson et al. 1989). Another possibility for overdispersion is intensifications of competitive interactions. Biotic interactions often intensify in response to resource availability, leading to competitive exclusion among closely related species (Webb 2000; Cavender-Bares et al. 2004; Cadotte et al. 2019). Considering that water serves as key resource for which mosses compete (Zamfir and Goldberg 2000), the overdispersion on wet barks could indicate a dominance of competition as a community assembly process mediated by resource availability.

Contrary to mosses, liverwort communities showed decreasing MPD and a tendency toward phylogenetic clustering (NRI < 0) in relation to the increased roughness, acidity, and wetness of barks (Fig. 2g, 2k). According to Fiz-Palacios et al. (2011), liverworts experienced a relatively slow diversification process from the mid-Cretaceous to the early Cenozoic era, during which mosses and ferns rapidly diversified in habitats created by angiosperms (as proposed by the “shadow of angiosperms” hypothesis; Schneider et al. 2004). It is possible that liverwort species, due to this constrained niche evolution, have maintained their specific habitats over time, leading to phylogenetic niche conservatism. The combination of closely related species having similar habitat preferences and environmental filtering associated with rough, acidic, and wet barks could have contributed to the observed pattern of phylogenetic clustering (Fig. 2k).

Ample evidence has shown that excess nitrogen owing to human activities (e.g., fertilization and atmospheric deposition) can reduce bryophyte richness (Oishi and Hiura 2017), both directly by posing toxic impacts and indirectly by enhancing the competitiveness of vascular plants (Turetsky 2003). In contrast, we found increasing species richness of both mosses and liverworts in response to inorganic nitrogen content (Fig. 2b, 2d), as represented by the second principal component (PC 2) (Fig. 1). The observed pattern could be attributed to the fact that our study was conducted in a pristine temperate forest where anthropogenic nitrogen inputs are kept minimal and epiphytic vascular plants are rare. In a nitrogen-limited environment with few competitors like our study site, barks with a high nitrogen content may serve as a hotspot for bryophytes. Nevertheless, it should be noted that we observed low levels of MPD on nitrogen-rich barks (Fig. 2f), indicating that only a restricted number of moss lineages could utilize such habitats.

In this study, we found that epiphytic bryophyte communities assemble non-randomly along gradients of bark traits (Fig. 2). Our study provides an important step toward understanding how host trees, as living patches, determine epiphyte assembly processes. Trees with different bark traits respond differently to environments (Rosell and Olson 2014), implying that potential future changes in tree bark diversity under environmental change can have cascading effects on epiphytic bryophyte diversity. While our study was based on snapshot data, future research should incorporate long-term monitoring and investigate the dynamics of host trees and epiphytes over time. Doing so would provide a more comprehensive understanding of epiphyte community assembly, which is essential for informing effective strategies for their conservation in the face of changing environments.

## Acknowledgements

We thank Takeshi Saeki and Takayuki Sugimoto for their assistance in the field. We are also grateful to Ryo Kitagawa and Xingfeng Si for their advice on statistical analyses, Yume Imada for her assistance during our pilot survey, Kentaro Fukushima for his advice on chemical analyses, and Yuta Kobayashi for his assistance in making figures.

## Authors contributions

ST conceived and led the study. All authors conducted the field survey. TO identified the bryophyte species. ST and WAA measured the bark traits. ST analysed the data. ST wrote the manuscript with inputs from other authors.

## Data availability

Bryophyte community matrix (presence-absence of 91 bryophyte species on 150 trees) is available at FigShare (https://doi.org/10.6084/m9.figshare.23673258).

## References

Bryant JA, Lamanna C, Morlon H, Kerkhoff AJ, Enquist BJ, Green JL (2008) Microbes on mountainsides: Contrasting elevational patterns of bacterial and plant diversity. Proc Natl Acad Sci 105:11505–11511. https://doi.org/10.1073/pnas.0801920105

Burns KC, Zotz G (2010) A hierarchical framework for investigating epiphyte assemblages: Networks, meta-communities, and scale. Ecology 91:377–385. https://doi.org/10.1890/08-2004.1

Cadotte MW, Carboni M, Si X, Tatsumi S (2019) Do traits and phylogeny support congruent community diversity patterns and assembly inferences? J Ecol 2065–2077. https://doi.org/10.1111/1365-2745.13247

Cadotte MW, Tucker CM (2017) Should environmental filtering be abandoned? Trends Ecol Evol 32:429–437. https://doi.org/10.1016/j.tree.2017.03.004

Callaway RM, Reinhart KO, Moore GW, Moore DJ, Pennings SC (2002) Epiphyte host preferences and host traits: Mechanisms for species-specific interactions. Oecologia 132:221–230. https://doi.org/10.1007/s00442-002-0943-3

Cavender-Bares J, Ackerly DD, Baum DA, Bazzaz FA (2004) Phylogenetic overdispersion in Floridian oak communities. Am Nat 163:823–843. https://doi.org/10.1086/386375

Ellis CJ (2012) Lichen epiphyte diversity: A species, community and trait-based review. Perspect Plant Ecol Evol Syst 14:131–152. https://doi.org/10.1016/j.ppees.2011.10.001

Fiz-Palacios O, Schneider H, Heinrichs J, Savolainen V (2011) Diversification of land plants: insights from a family-level phylogenetic analysis. BMC Evol Biol 11:341. https://doi.org/10.1186/1471-2148-11-341

Gerhold P, Cahill JF, Winter M, Bartish I V., Prinzing A (2015) Phylogenetic patterns are not proxies of community assembly mechanisms (they are far better). Funct Ecol 29:600–614. https://doi.org/10.1111/1365-2435.12425

Gotelli NJ (2000) Null model analysis of species co-occurrence patterns. Ecology 81:2606–2621. https://doi.org/10.2307/177478

Gustafsson L, Eriksson I (1995) Factors of Importance for the Epiphytic Vegetation of Aspen Populus tremula with Special Emphasis on Bark Chemistry and Soil Chemistry. J Appl Ecol 32:412. https://doi.org/10.2307/2405107

Kaufmann S, Weinrich T, Hauck M, Leuschner C (2019) Vertical variation in epiphytic cryptogam species richness and composition in a primeval Fagus sylvatica forest. J Veg Sci 30:881–892. https://doi.org/10.1111/jvs.12775

Lamit LJ, Lau MK, Næsborg RR, Wojtowicz T, Whitham TG, Gehring CA (2015) Genotype variation in bark texture drives lichen community assembly across multiple environments. Ecology 96:960–971. https://doi.org/10.1890/14-1007.1

Löbel S, Rydin H (2010) Trade-offs and habitat constraints in the establishment of epiphytic bryophytes. Funct Ecol 24:887–897. https://doi.org/10.1111/j.1365-2435.2010.01705.x

Mendieta-Leiva G, Zotz G (2015) A conceptual framework for the analysis of vascular epiphyte assemblages. Perspect Plant Ecol Evol Syst 17:510–521. https://doi.org/10.1016/j.ppees.2015.09.003

Mitchell RJ, Hewison RL, Beaton J, Douglass JR (2021) Identifying substitute host tree species for epiphytes: The relative importance of tree size and species, bark and site characteristics. Appl Veg Sci 24:1–13. https://doi.org/10.1111/avsc.12569

Oishi Y, Hiura T (2017) Bryophytes as bioindicators of the atmospheric environment in urban-forest landscapes. Landsc Urban Plan 167:348–355. https://doi.org/10.1016/j.landurbplan.2017.07.010

Robinson AL, Vitt DH, Timoney KP (1989) Patterns of Community Structure and Morphology of Bryophytes and Lichens Relative to Edaphic Gradients in the Subarctic Forest-Tundra of Northwestern Canada. Bryologist 92:495–512. https://doi.org/10.2307/3243674

Rosell JA, Olson ME (2014) The evolution of bark mechanics and storage across habitats in a clade of tropical trees. Am J Bot 101:764–777. https://doi.org/10.3732/ajb.1400109

Schneider H, Schuettpelz E, Pryer KM, Cranfill R, Magallón S, Lupia R (2004) Ferns diversified in the shadow of angiosperms. Nature 428:553–557. https://doi.org/10.1038/nature02361

Shi S, Huang Y, Zeng K, Tan F, He H, Huang J, Fu Y (2005) Molecular phylogenetic analysis of mangroves: Independent evolutionary origins of vivipary and salt secretion. Mol Phylogenet Evol 34:159–166. https://doi.org/10.1016/j.ympev.2004.09.002

Spicer ME, Woods CL (2022) A case for studying biotic interactions in epiphyte ecology and evolution. Perspect Plant Ecol Evol Syst 54:125658. https://doi.org/10.1016/j.ppees.2021.125658

Stech M, Quandt D (2010) 20,000 species and five key markers: The status of molecular bryophyte phylogenetics. Phytotaxa 9:196–228. https://doi.org/10.11646/phytotaxa.9.1.11

Tatsumi S, Cadotte MW, Mori AS (2019) Individual-based models of community assembly: Neighbourhood competition drives phylogenetic community structure. J Ecol 107:735–746. https://doi.org/10.1111/1365-2745.13074

Tatsumi S, Ohgue T, Azuma W, Tuovinen V, Imada Y, Mori AS, Thor G, Ranlund Å (2017) Tree hollows can affect epiphyte species composition. Ecol Res 32. https://doi.org/10.1007/s11284-017-1468-x

Turetsky MR (2003) New frontiers in bryology and lichenology: The role of bryophytes in carbon and nitrogen cycling. Bryologist 106:395–409. https://doi.org/10.1639/05

Webb CO (2000) Exploring the phylogenetic structure of ecological communities: An example for rain forest trees. Am Nat 156:145–155. https://doi.org/10.1086/303378

Webb CO, Ackerly DD, McPeek MA, Donoghue MJ (2002) Phylogenies and community ecology. Annu Rev Ecol Syst 33:475–505. https://doi.org/10.1146/annurev.ecolsys.33.010802.150448

Wyse S V., Burns BR (2011) Do host bark traits influence trunk epiphyte communities? N Z J Ecol 35:296–301

Zamfir M, Goldberg DE (2000) The effect of initial density on interactions between bryophytes at individual and community levels. J Ecol 88:243–255. https://doi.org/10.1046/j.1365-2745.2000.00442.x

