## Supplementary materials for "Bark traits affect epiphytic bryophyte community assembly in a temperate forest"

**Supplementary text 1: *Bryophyte phylogeny reconstruction***

A bryophyte phylogeny was reconstructed based on three chloroplast genes (*rbc*L, *rps*4, and *trn*L-F) that are commonly used in bryophyte phylogenetics (Stech and Quandt 2010). Sequence data were retrieved from GenBank. *Chara* spp. were included as outgroup species representing the early divergence of Streptophyta lineages. For species of which the sequences for *rbc*L and/or *rps*4 were unavailable, the sequences were substituted by those of the congeneric species whenever possible. Sequences were aligned using the E-INS-i algorithm in MAFFT 7 (Katoh and Standley 2013) separately for each gene. The best substitution model was selected based on the corrected Akaike information criterion using MEGA 7 (Kumar et al. 2016), which resulted in the GTR+Г+I model being selected for all genes. The posterior distributions of the model parameters were estimated using the Markov chain Monte Carlo method implemented by MrBayes 3.2 (Ronquist et al. 2012). The phylogram was transformed to a chronogram using the semiparametric rate smoothing based on penalized likelihood (Sanderson 2002) with smoothing parameter λ = 39.8. The chronogram was calibrated with node ages *sensu* Cooper et al. (2012).

**Reference**

Cooper ED, Henwood MJ, Brown EA (2012) Are the liverworts really that old? Cretaceous origins and Cenozoic diversifications in Lepidoziaceae reflect a recurrent theme in liverwort evolution. Biol J Linn Soc 107:425–441. https://doi.org/10.1111/j.1095-8312.2012.01946.x

Katoh K, Standley DM (2013) MAFFT multiple sequence alignment software version 7: Improvements in performance and usability. Mol Biol Evol 30:772–780. https://doi.org/10.1093/molbev/mst010

Kumar S, Stecher G, Tamura K (2016) MEGA7: Molecular evolutionary genetics analysis version 7.0 for bigger datasets. Mol Biol Evol 33:1870–1874. https://doi.org/10.1093/molbev/msw054

Ronquist F, Teslenko M, Van Der Mark P, Ayres DL, Darling A, Höhna S, Larget B, Liu L, Suchard MA, Huelsenbeck JP (2012) Mrbayes 3.2: Efficient bayesian phylogenetic inference and model choice across a large model space. Syst Biol 61:539–542. https://doi.org/10.1093/sysbio/sys029

Sanderson MJ (2002) Estimating absolute rates of molecular evolution and divergence times: A penalized likelihood approach. Mol Biol Evol 19:101–109. https://doi.org/10.1093/oxfordjournals.molbev.a003974

Stech M, Quandt D (2010) 20,000 species and five key markers: The status of molecular bryophyte phylogenetics. Phytotaxa 9:196–228. https://doi.org/10.11646/phytotaxa.9.1.11

**Supplementary text 2: *Bark trait measurement***

We measured bark roughness, water content, pH, and inorganic nitrogen content (NH_4_^+^, NO_2_^−^, and NO_3_^−^) for the 10 studied tree species. These traits were selected based on previous findings that they often show significant associations with epiphyte community structure (Gustafsson & Eriksson 1995; reviewed in Ellis 2012). Bark roughness was defined as the arc length of the stem, following the idea of Glitzenstein and Harcombe (1979). We horizontally oriented a 10-cm adjustable-curve ruler that does not penetrate into crevices of the bark, and measured the distance between the stem surface and the ruler at 0.5-cm intervals (i.e., 21 points) along the ruler. For each tree species, the bark roughness was measured at three random points (at 50–200 cm height) on each of the three trees which were randomly selected in the field. For rest of the traits, bark samples were collected at the same points where the roughness was measured. We used the outer bark (i.e., rhytidomes) for measurements, after shearing off the vascular cambiums and xylems from the samples in the laboratory. Bark water content was measured by weighing the samples before and after drying. Bark pH and nitrogen content were measured by reference to Gustafsson and Eriksson (1995). Specifically, the bark samples were pooled for each tree individual, chipped into ca. 5-mm squares, and shook in Milli-Q water (undried bark = 3–5 g, bark:water = 1:10 w/v) at 200 rpm for 1 h. The pH of the solutions was then measured with an electrode (D-53, Horiba, Kyoto, Japan). For the inorganic nitrogen content, the solutions were further syringe-filtered and measured with a colorimetric method using an auto-analyzer (AutoAnalyzerIII, BL TEC, Osaka, Japan). The species’ mean trait values were used for statistical analyses.

**References**

Ellis CJ (2012) Lichen epiphyte diversity: A species, community and trait-based review. Perspect Plant Ecol Evol Syst 14:131–152. https://doi.org/10.1016/j.ppees.2011.10.001

Glitzenstein J, Harcombe PA (1979) Site-specific changes in the bark texture of *Quercus falcata* Michx. (Southern Red Oak). Am J Bot 66:668–672. https://doi.org/10.2307/2442411

Gustafsson L, Eriksson I (1995) Factors of importance for the epiphytic vegetation of aspen *Populus tremula* with special emphasis on bark chemistry and soil chemistry. J Appl Ecol 32:412. https://doi.org/10.2307/2405107

**Table S1.** Bark trait values of ten tree species.

| Tree species | Roughness (mm)^†^ | Water content (%) | pH | Inorganic nitrogen content (μg/g) ^‡^ |
| --- | --- | --- | --- | --- |
| *Acer pictum* | 0.701 | 54.9 | 6.15 | 1.09 |
| *Acer sieboldianum* | 0.704 | 46.8 | 5.45 | 0.37 |
| *Aesculus turbinata* | 1.021 | 207.5 | 5.60 | 0.09 |
| *Betula grossa* | 0.855 | 56.1 | 4.99 | 11.04 |
| *Castanea crenata* | 3.178 | 155.6 | 4.51 | 4.67 |
| *Clethra barbinervis* | 1.064 | 112.1 | 5.16 | 4.11 |
| *Cryptomeria japonica* | 2.847 | 115.6 | 4.16 | 1.64 |
| *Fagus crenata* | 0.742 | 55.8 | 6.18 | 5.67 |
| *Quercus crispula* | 1.367 | 164.0 | 5.55 | 15.43 |
| *Quercus serrata* | 3.212 | 106.6 | 4.52 | 3.47 |

^†^ Measured as the arc length of the stem minus the length of adjustable-curve ruler (100 mm). The larger the values are, the rougher the bark is.

^‡^ Measured as the amount of ion mass (NH_4_^+^, NO_2_^−^, and NO_3_^−^; μg) in extracted solutions per 1 g of bark.

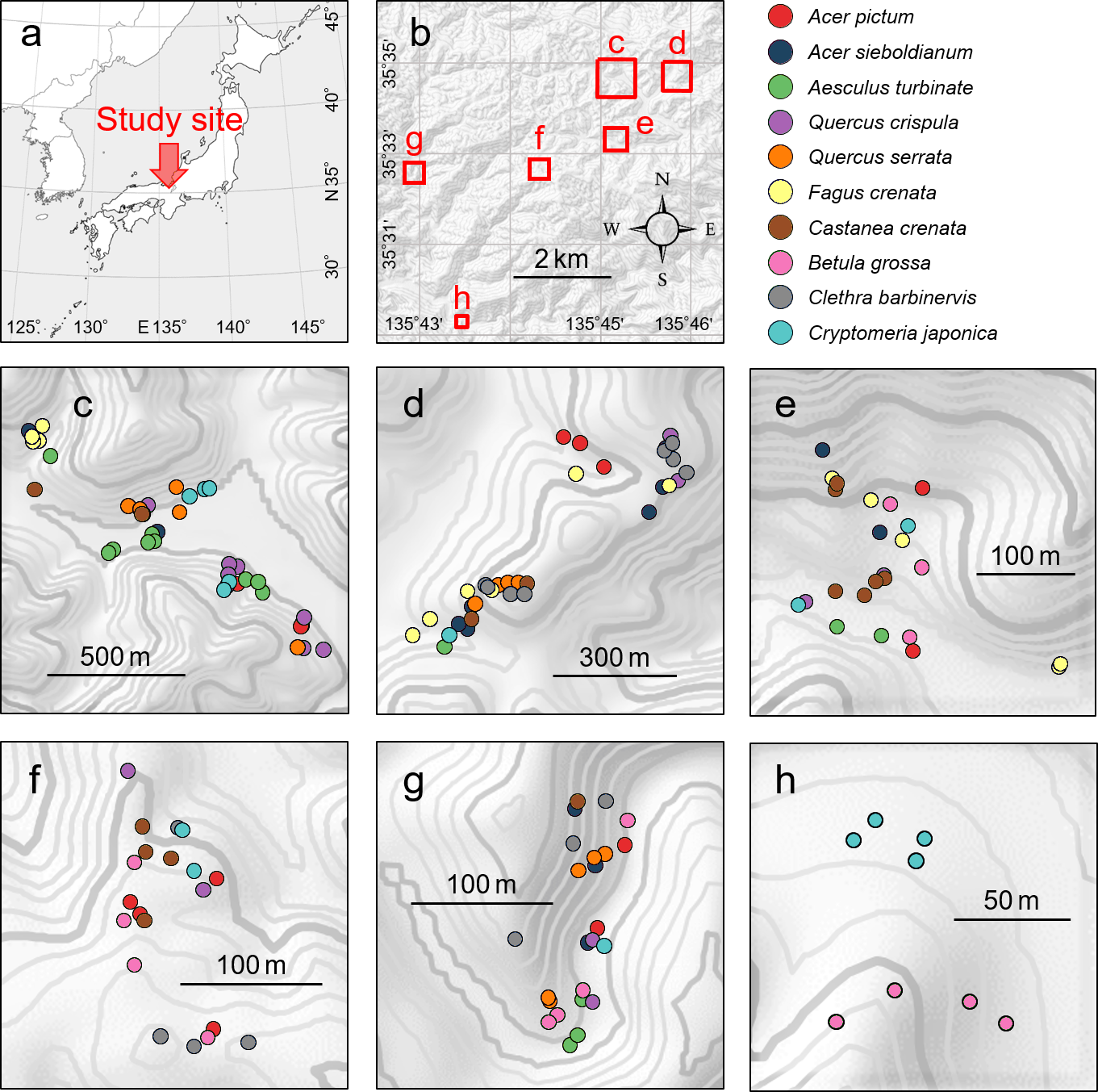

**Figure S1.** Locations of the (a) study site, (b) plots, and (c–h) trees. The letters “c–h” in panel b corresponds to the locations of plots shown in panels c–h. The thin and thick lines in panels c–h indicate 10-m and 50-m contours, respectively.

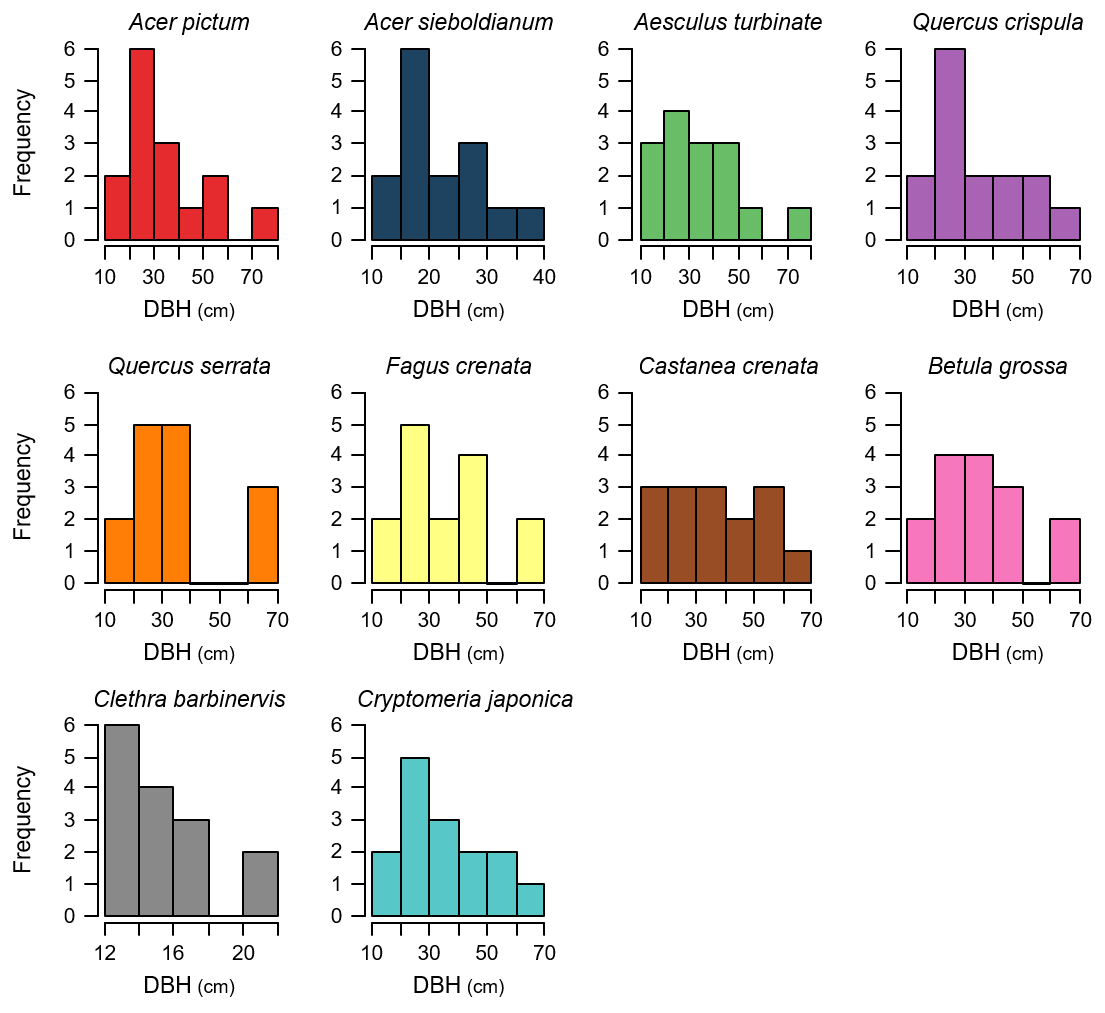

**Figure S2.** Frequencies of diameter at breast height (DBH) of the studied species.
